# Restricted differentiative capacity of Wt1-expressing peritoneal mesothelium in postnatal and adult mice

**DOI:** 10.1101/2020.08.21.257154

**Authors:** Thomas P Wilm, Helen Tanton, Fiona Mutter, Veronica Foisor, Ben Middlehurst, Kelly Ward, Tarek Benameur, Nicholas Hastie, Bettina Wilm

## Abstract

Previously, genetic lineage tracing based on the mesothelial marker Wt1, appeared to show that peritoneal mesothelial cells have a range of differentiative capacities and are the direct progenitors of vascular smooth muscle in the intestine. However, it was not clear whether this was a temporally limited process or continued throughout postnatal life. Here, using a conditional Wt1-based genetic lineage tracing approach, we demonstrate that the postnatal and adult peritoneum covering intestine, mesentery and body wall only maintained itself and failed to contribute to other visceral tissues. Pulse-chase experiments of up to 6 months revealed that Wt1-expressing cells remained confined to the peritoneum and failed to differentiate into cellular components of blood vessels or other tissues underlying the peritoneum. Ablation of Wt1 in adult mice did not result in changes to the intestinal wall architecture. In the heart, we observed that Wt1-expressing cells maintained the epicardium and contributed to coronary vessels in newborn and adult mice. Our results demonstrate that Wt1-expressing cells in the peritoneum have limited differentiation capacities, and that contribution of Wt1-expressing cells to cardiac vasculature is based on organ-specific mechanisms.

## Introduction

The mesothelium of parietal and visceral peritoneum arises as part of the serosa of the peritoneal cavity from the lateral plate mesoderm, where it forms a simple squamous epithelium covering all organs (visceral) and the body wall musculature (parietal) within the cavity ^1,2^. The Wilms’ tumour protein 1 (Wt1) is a key marker of mesothelia, and a transgenic mouse line expressing Cre recombinase under control of regulatory elements of the human *WT1* gene (Tg(WT1-cre)AG11Dbdr, in short: Wt1-Cre) had been previously used to track mesothelial cells in mice ^1^. In adult Wt1-Cre; Rosa26^LacZ/LacZ^ compound mutant mice, XGal-labelled vascular smooth muscle cells (VSMCs) were found in the vasculature of the mesentery and intestine, as well as the heart and lungs ^1,3^. These findings led us to conclude that cells of the visceral mesothelium give rise directly to VSMCs in the intestine and mesentery ^1^. Studies using a different Wt1-Cre line (Tg(Wt1-cre)#Jbeb) revealed a similar contribution of Wt1-expressing cells to the mesothelium and the visceral and vascular smooth muscle of the developing intestine ^4^.

However, these genetic lineage tracing systems were unable to distinguish the time window during which cells had expressed Wt1, as once tagged, the cells were consequently irreversibly labelled. Therefore, it remains unclear whether Wt1-expressing mesothelial cells give rise to VSMCs continuously throughout life. Furthermore, it is not clear whether postnatal or even adult mesothelial cells in the peritoneal cavity generally contribute to the maintenance of the intestinal and parietal body wall. This is pertinent because of reports suggesting that mesothelial cells have the plasticity to differentiate into cells of different mesodermal lineages ^5–8^.

To address these questions, we determined the spatio-temporal contribution of Wt1-expressing cells to the adult and postnatal intestine and peritoneal body wall, including the mesentery. For this purpose, we used a conditional Wt1-driven Cre system (Wt1^tm2(cre/ERT2)Wtp^) which relies on tamoxifen administration to activate recombination and thus lineage tracing ^9,10^. Our comparison of short- and long-term lineage pulse-chase experiments revealed that Wt1-lineage traced mesothelial cells in neonatal or adult visceral and parietal peritoneum only contributed to maintenance of cells of the serosa (and mesenteric fat) and not to vascular cells or intestinal wall tissue. This was different in the heart, where cells of the coronary vasculature were labeled already after a short pulse-chase period, indicating postnatal contribution of Wt1-expressing cells. In addition, loss of Wt1 in adult visceral mesothelium failed to interfere with the overall tissue integrity of the intestinal wall. Our findings clearly demonstrate the specific and limited lineage of Wt1-expressing mesothelial cells of the peritoneum in healthy postnatal and adult mice.

## Results

### Adult Wt1-derived cells maintain the visceral and parietal mesothelium without contribution to vascular smooth muscle

We examined whether mesothelial cells (MCs) contribute to the formation of vascular cells and other tissue components in the adult intestine. After a 2-4 week chase period in Wt1^CreERT2/+^; Rosa26^LacZ/LacZ^ and Wt1^CreERT2/+^; Rosa26^mTmG/mTmG^ mice we found lineage-traced cells in a patchy pattern in the intestinal and parietal serosa (Figure 1A, B). We could not detect any contribution of labelled Wt1-expressing mesothelial cells to vascular or non-vascular smooth muscle in the intestine (Figure 1C, D), or other cell types of body or intestinal wall, or the mesentery, except for the previously reported contribution to mesenteric adipose cells, which was confirmed by histological analysis (Supplementary Figure S1A, B) ^11,12^. However, in the visceral (mesentery and omentum) and parietal peritoneum, we observed a recently described Wt1-expressing submesothelial mesenchymal population (Figure 2A-D)^13^.

**Figure 1:**
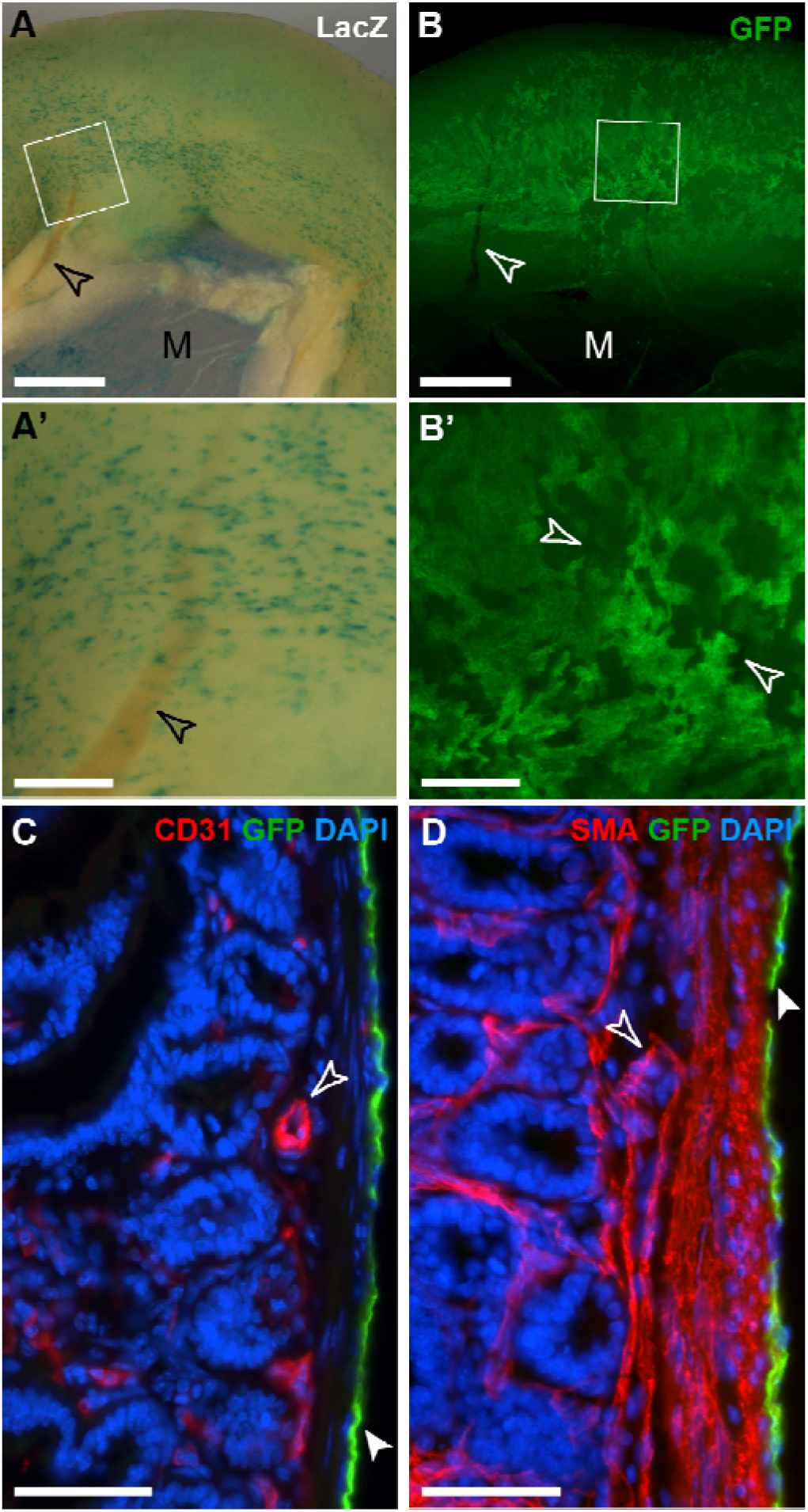
Whole mount and frozen section analysis of adult intestine after lineage tracing of Wt1-expressing cells. Adult mice with either the Wt1^CreERT2/+^; Rosa26^LacZ/LacZ^ (A) or the Wt1^CreERT2/+^; Rosa26^mTmG/mTmG^ (B-D) reporter system were analysed 2-4 weeks after Tamoxifen administration. A, B. Labelled cells were found on the surface of the intestine and in the mesentery (M) in a random and patchy distribution, but no contribution of labelled cells to the vascular or visceral smooth muscle of the intestine or mesentery was observed (open arrowheads; A’, B’, magnifications of highlighted areas). C, D. Immunofluorescence labelling of frozen sections from Wt1^CreERT2/+^; Rosa26^mTmG/mTmG^ intestine. Wt1-lineage traced cells were solely identified in the mesothelium (filled arrowheads; open arrowheads pointing to vascular tissue labelled with endothelial CD31 and α-smooth muscle actin (SMA) antibodies). Scale bars, 1 mm (A, B), 200 μm (A’, B’), 50 μm (C, D).

**Figure 2:**
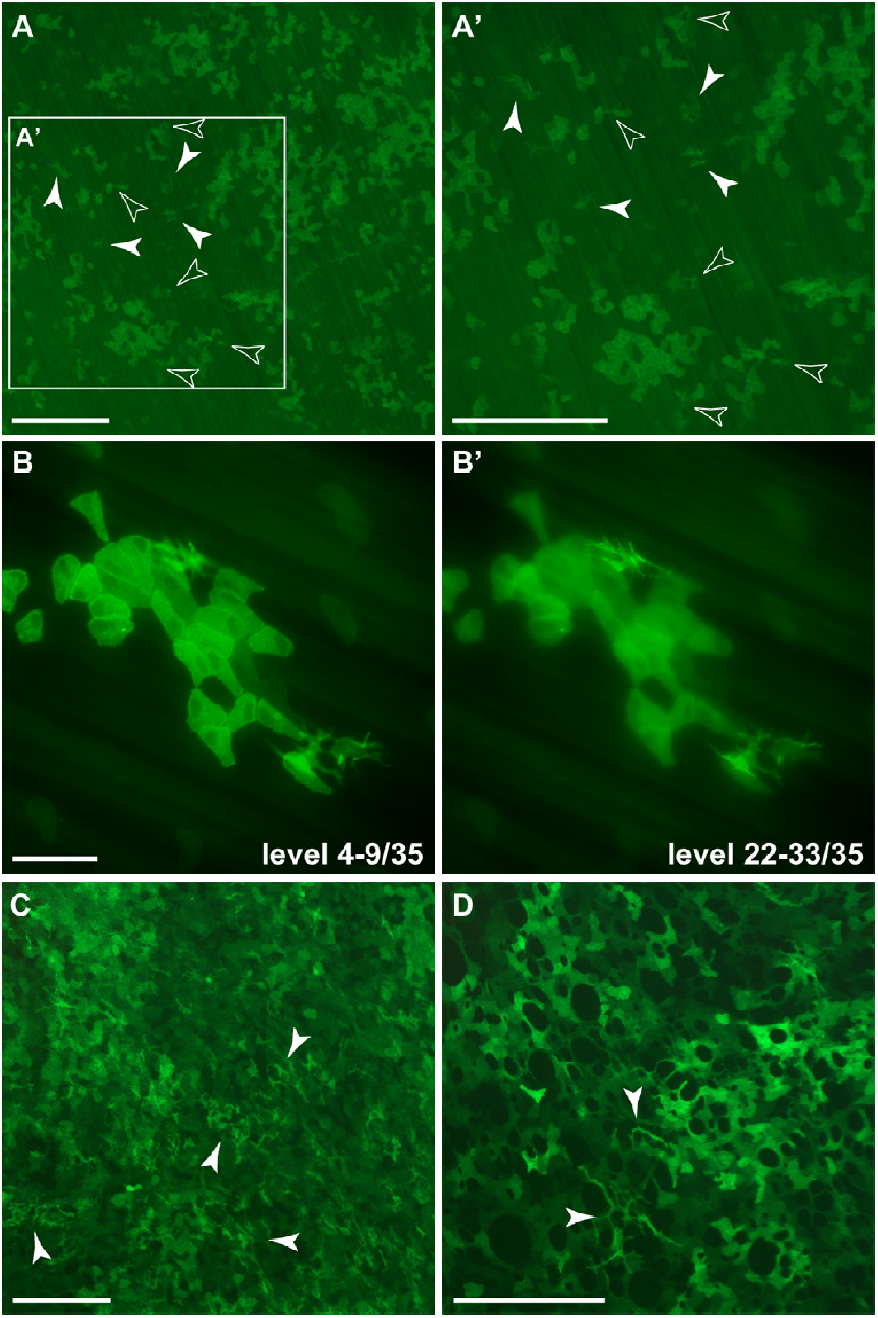
Lineage tracing of Wt1-expression GFP-positive mesenchymal cells in parietal and visceral peritoneum. Adult mice with the Wt1^CreERT2/+^; Rosa26^mTmG/mTmG^ reporter system were analysed 2-4 weeks after tamoxifen administration. A, A’. In the parietal peritoneum of the body wall, GFP-positive mesothelial cells were detected in single and clustered patterns. GFP-positive mesenchymal cells were found in areas without GFP-positive mesothelial cells (solid arrow heads) as well as close to or underneath GFP-positive mesothelial clusters (transparent arrow heads). B, B’. Partial projections of a 0.5 μm distanced 35 focal level Z-stack; levels 4-9 (B) show a cluster of GFP-positive mesothelial cells and levels 22-33 (B’) three GFP-positive individual mesenchymal cells located underneath the mesothelial layer. C, D. GFP-positive mesenchymal cells (arrowheads) scattered between GFP+ mesothelial cells in the visceral peritoneum of the mesentery (C) and the omentum (D). Scale bars, 500 μm (A, A’, C, D), 200 μm (B, B’).

We confirmed by direct comparison of GFP and XGal staining in the intestinal and parietal serosa from compound mutant Wt1^CreERT2/+^; Rosa26^LacZ/mTmG^ mice that both reporter systems gave very similar results (Supplementary Figure S2). It is important to note, however, that GFP^+^ cells were not necessarily LacZ^+^ and vice versa, although overall there appeared to be a larger number of GFP^+^ cells present compared to LacZ^+^, and their signal was more clearly detectable.

To determine whether the previously reported contribution of MCs to the intestinal vascular smooth muscle compartment is controlled by a slow process of tissue homeostasis ^1^, we performed a long-term chase experiment for 6 months. Our data revealed that even 6 months after the tamoxifen pulse, coverage of the visceral mesothelium with labelled cells was still patchy, and no contribution to intestinal vascular smooth muscle or any other intestinal wall cells could be detected (Supplementary Figure S3). To probe this further, we used the Wt1^CreERT2/+^; Rosa26^Confetti/+^ reporter system which allows the detection of clonal expansion in lineage tracing studies. When comparing labelled cells in the intestinal serosa in mice after 2 weeks or 6 months pulse-chase experiments, we observed a limited clonal expansion of labelled visceral MCs in adult mice over time (Figure 3A, B). While there were significantly more mesothelial 2-cell clones in the serosa after 6 months, clones that contained 3 or more cells had not significantly increased after the long chase-period (Figure 3C, D). Taken together, our observations indicate that Wt1-derived MCs in the peritoneum of healthy adult mice were restricted to maintaining its homeostasis, possibly at a low turnover rate ^14^.

**Figure 3:**
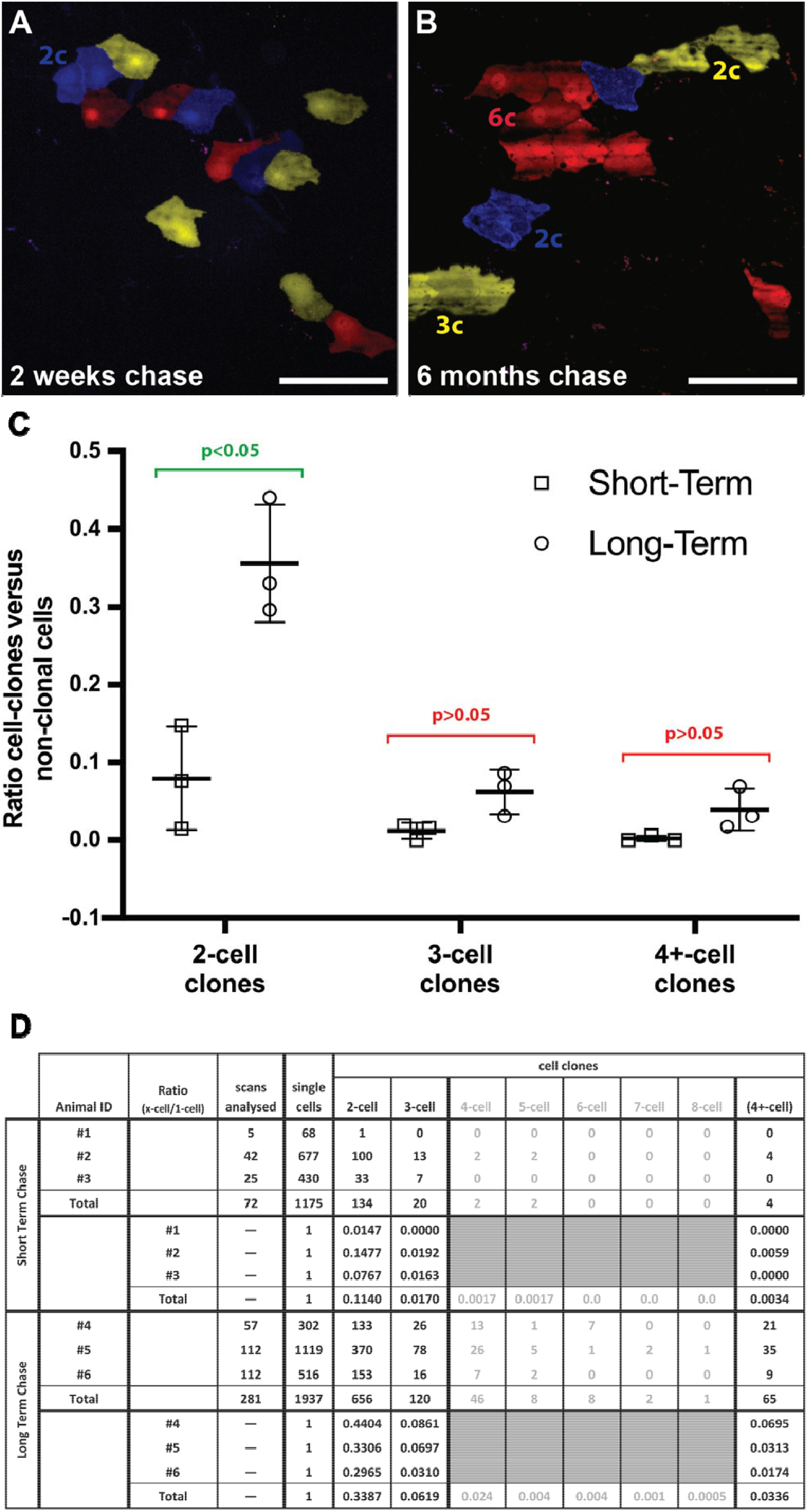
Clonal analysis of short-term and long-term lineage tracing of Wt1-expressing cells in adult mouse intestine. Adult mice with the Wt1^CreERT2/+^; Rosa26^cofetti/LacZ^ reporter system were analysed 2 weeks and 6 months after Tamoxifen administration, respectively. Random segments (8 per animal; group size n = 3) of intestine were analysed by confocal microscopy after short fixation to eliminate peristalsis. Random Z-stacks were recorded and the number of cells scored into two groups: individual cells for all three reporters (as defined by no contact to same-colour cells) and cell clones (as defined by contact to same-colour cells). A, B. Exemplary images of intestine segments from short-term (A) and long-term chase experiments (B). The image of the short-term experiment shows one 2-cell clone (blue, 2c), while that of the long-term experiment shows one 6-cell clone (red, 6c), one 3-cell clone (yellow, 3c) and two 2-cell clones (blue and yellow, 2c). Individual non-clonal cells were found as well. C. Grouped Scatter Graph showing the ratios of cell-clone numbers versus non-clonal cell numbers from individual animals including the standard deviation of the mean (group size n = 3). Statistical analysis was performed using unpaired multiple t-tests with Holm-Šídák multiple comparisons correction for 2-cell- (adjusted p value = 0.026), 3-cell- and 4+-cell-clone ratios (both adjusted p value = 0.086). D. Table summarizing the scores for clonal and non-clonal cells including ratio for individual animals of short- and long-term chase and pooled totals. Scale bars are 100 μm (A, B).

### Wt1 expression is not required for the maintenance of the visceral serosa and the intestinal wall

Because Wt1 is expressed in the adult visceral and parietal peritoneum, we asked whether it is required for maintenance of the serosa and underlying tissues. To address this question, we ablated Wt1 in adult mesothelial cells by using Wt1^CreERT2/co^ mice, similar to the approach described by Chau and colleagues ^12^. Because the Wt1^CreERT2/+^ mutation leads to a loss of Wt1 expression on this allele, administration of tamoxifen in Wt1^CreERT2/co^ mice resulted in loss of Wt1 on both alleles simultaneously. In adult mice with Wt1 ablation, we observed loss of mesenteric fat as one of the characteristics previously described (not shown) ^12^. In addition, the health in all animals lacking Wt1 severely deteriorated from approximately day 10 after the start of tamoxifen administration onwards ^12^. This precluded a detailed analysis of the longer-term impact of loss of Wt1 on the homeostasis of the peritoneum and in particular the visceral lining of the intestine and its underlying layers. However, our histological analysis of various regions of the intestine at day 10 after tamoxifen administration showed no effect on the presence and appearance of the mesothelium, or the overall morphology of the intestinal wall (Figure 4).

**Figure 4:**
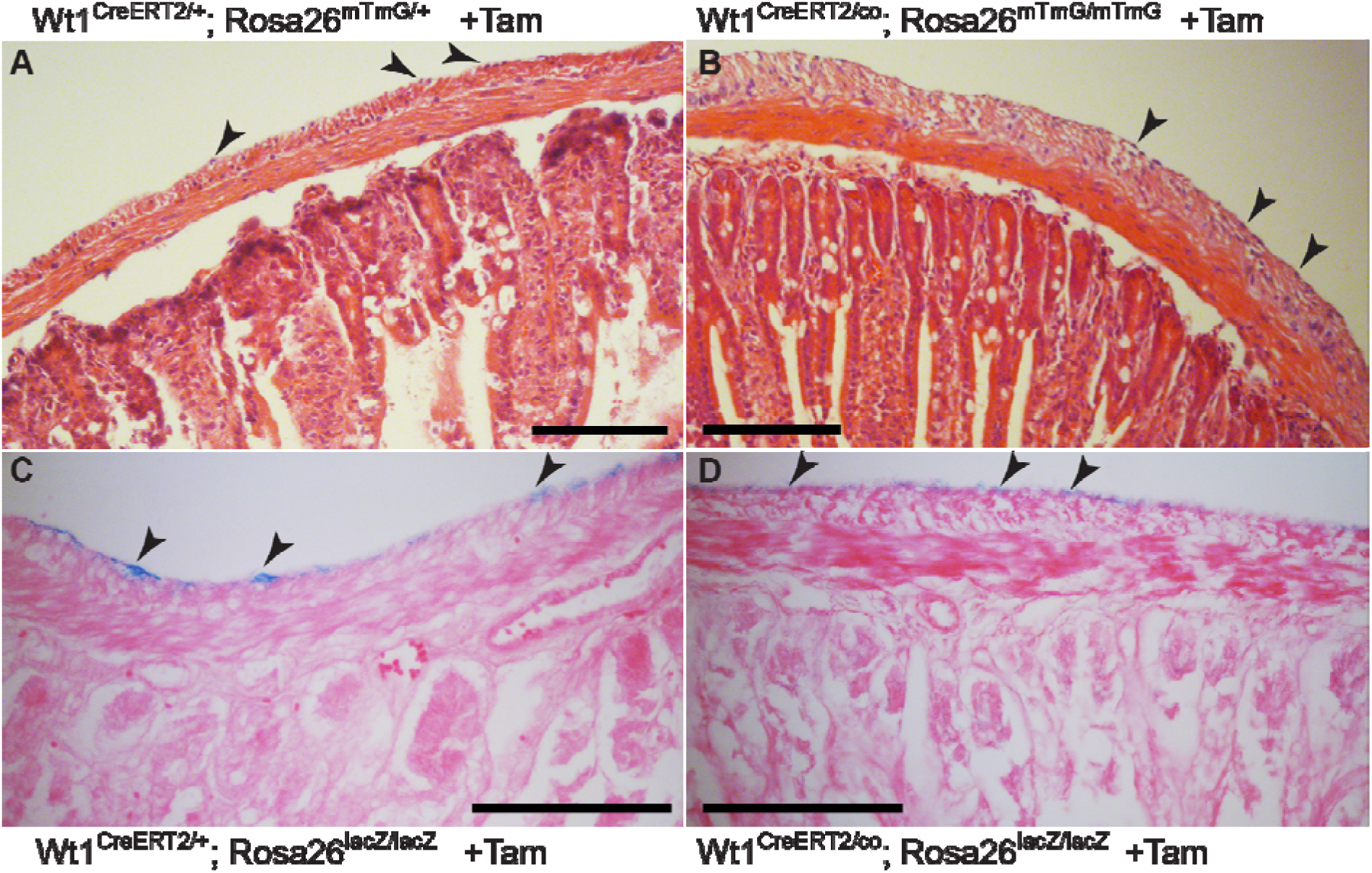
Ablation of Wt1 in adult intestine. Tamoxifen was given to mice carrying either the Wt1^CreERT2/co^ or the Wt1^CreERT2/+^ allele, for 5 consecutive days. Mice were sacrificed on day 10 and the intestines harvested and XGal stained and/or directly processed for histology (Haematoxylin & Eosin or Eosin counter stain). After Tamoxifen, mice with and without the conditional Wt1 allele showed normal architecture of the intestinal wall, with mesothelial cells present as visualised by nuclear Haematoxylin staining (A, B) or by XGal staining (C, D). Scale bars, 100 μm (A-D).

### Wt1-derived cells in the adult heart maintain the epicardium and contribute to coronary vessel cells

We analysed the contribution of Wt1-derived cells in the adult heart. Lineage-traced Wt1-derived epicardial cells were found in patches over the heart in adult mice (Figure 5A, B, Supplementary Figure S1C), but XGal- or GFP-labelled cells also formed part of the coronary vasculature of the heart even after a 2-week chase period (Figure 5A’, Supplementary Figure S1D). The coronary contribution was more difficult to determine in whole mount tissue analysis of mice carrying the Wt1^CreERT2/+^; Rosa26^mTmG/mTmG^ reporter because of the strong coverage of GFP-expressing cells in the epicardium (Figure 5B’). Immunohistological analysis of the heart wall showed that Wt1-derived GFP^+^ cells localised around coronary vessels and co-expressed the classical vascular smooth muscle cell marker α-smooth muscle actin (SMA), or the endothelial marker CD31 (Figure 5C, D). We also included kidneys in our analysis because adult podocytes express Wt1, thus allowing assessment of successful tamoxifen administration. Adult kidneys showed the expected labelling of glomeruli two weeks after tamoxifen administration (Figure 5E-H), where GFP-positive cells co-expressed Wt1, indicating their podocyte identity (Figure 5H’-H’’’). We occasionally (in 1 out of 10 analysed kidneys) observed one or two LacZ- or GFP-positive tubules in the medulla region of the sagittally dissected kidneys (Figure 5I, J), suggesting that there was a small population of Wt1-expressing cells in the adult kidney with the potential to contribute to the tubular elements of the nephrons.

**Figure 5:**
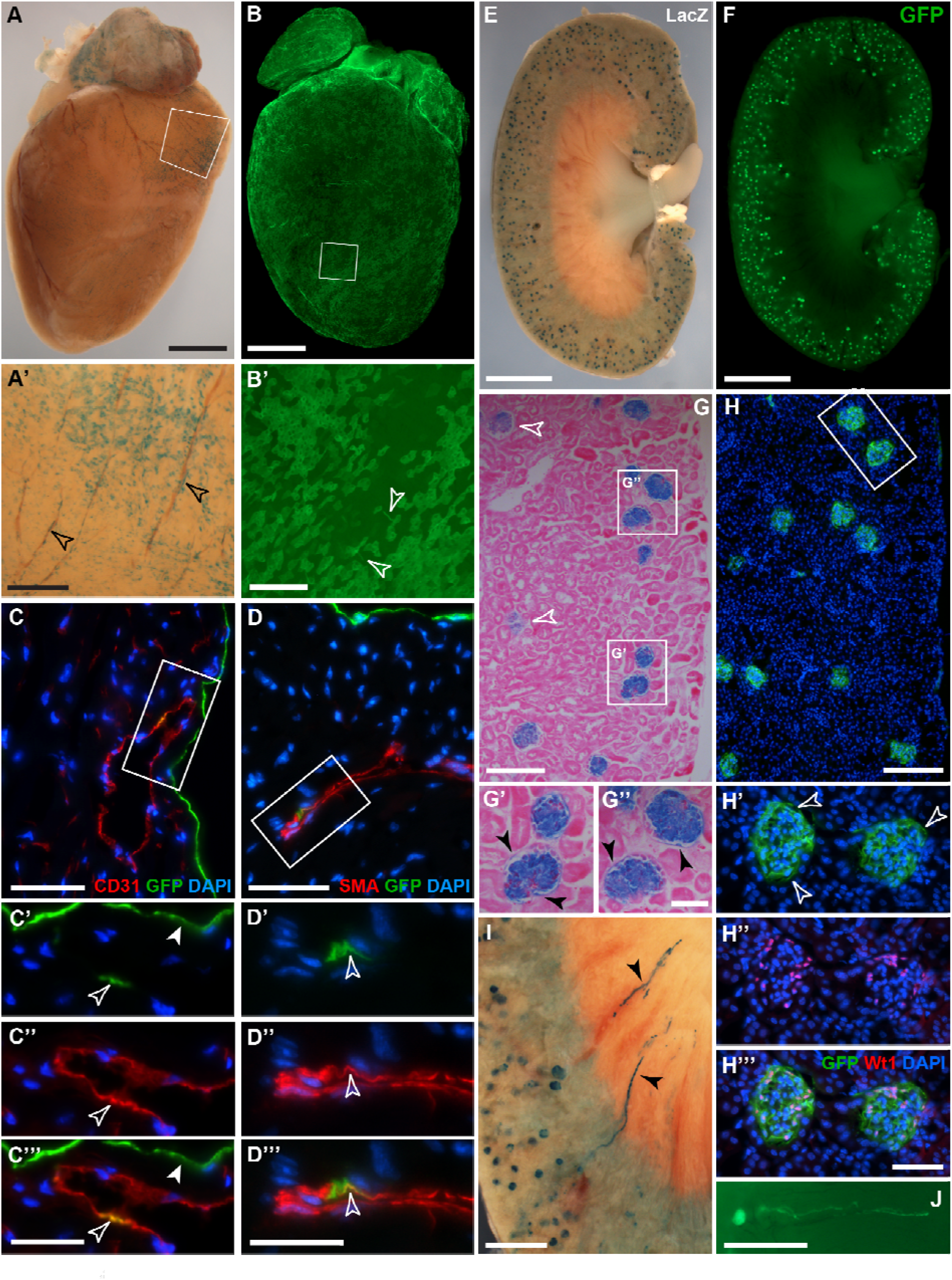
Whole mount and histological analysis of adult heart and kidney after lineage tracing of Wt1-expressing cells. Adult mice with either the Wt1^CreERT2/+^; Rosa26^LacZ/LacZ^ or the Wt1^CreERT2/+^; Rosa26^mTmG/mTmG^ reporter system were analysed 2-4 weeks after Tamoxifen administration. A, B. Coverage of the epicardium with labelled cells was patchy. A’, B’. Magnification of highlighted areas in A and B. LacZ-positive cells appeared to localize around coronary vessels (arrowheads) in addition to presence in the epicardium. By contrast, GFP-labelled cells consisted predominantly of flat-shaped epicardial cells, with a few thinly shaped cells detectable (open arrowheads). C, D. Immunofluorescence labelling of frozen sections from Wt1^CreERT2^; Rosa26^mTmG/mTmG^ heart tissue. Co-expression of GFP with CD31 or SMA, respectively, was observed in coronary vasculature. C’-C’’’, D’-D’’’. Magnification of areas highlighted in C and D. GFP and DAPI (C’, D’); CD31 or SMA and DAPI (C’’, D’’), merged (C’’’, D’’’; co-localization highlighted by arrow heads). E, F. In kidney whole mounts (sagittal halves), labelled cells expressing LacZ or GFP were found in the glomeruli. G. Eosin-counterstained sagittal paraffin sections showing LacZ-expression in the glomeruli of the kidney (open arrowheads pointing to glomeruli showing weaker XGal staining due to reduced penetration of staining reagents into tissue). G’, G’’. LacZ-expressing cells are also detected in the parietal epithelial layer of the Bowman Capsule (filled arrowheads). H. Immunofluorescence in frozen sagittal kidney sections revealed GFP-expressing cells in the glomeruli. H’-H’’’. GFP-labelled cells co-expressed Wt1 (H’, GFP and DAPI; H’’, Wt1 and DAPI; H’’’, merged). I, J. In rare cases, LacZ- or GFP-expressing cells were found in tubular structures reaching into the renal medulla (filled arrowheads). Scale bars, 1.5 mm (A, B), 400 μm (A’), 200 μm (B’), 50 μm (C, D), 25 μm (C’-C’’’, D’-D’’’), 2 mm (E, F), 300 μm (G, H), 100 μm (G’, G’’, H’-H’’’), 700 μm (I, J).

### Wt1-derived cells in newborn lineage tracing reveal wider contribution to the heart and kidneys, but not to the peritoneum

Since the adult mesothelium of the peritoneum appeared to be restricted to its own maintenance, we tested whether mesothelial cells may still have some degree of plasticity either in juvenile mice directly after weaning or within the first few days after birth ^15–18^, and thus would show capacity to contribute to intestinal tissue homeostasis or to differentiate into VSMCs. After tamoxifen administration in four weeks old juvenile Wt1^CreERT2/+^; Rosa26^LacZ/+^ mice and chase periods between 7 and 17 weeks, restricted contributions of Wt1-derived LacZ-positive cells were found in similar locations to those in adult mice (data not shown). Correspondingly, after initiating a 7-weeks chase period in newborn Wt1^CreERT2/+^; Rosa26^LacZ/+^ mice, we detected coverage of LacZ-positive cells in the visceral mesothelium comparable to adult mice (Figure 6A), and no contribution to the vasculature of the mesentery or intestine, or within the intestinal wall (Figure 6B, C). By contrast, lineage-traced cells were found as expected in the epicardial layer of the heart and also contributing to its coronary and micro-vasculature (Figure 6D-F). These results indicate that while in the newborn, Wt1-expressing epicardial cells gave rise to coronary vessels in line with reported plasticity of the newborn heart ^17,19–21^, visceral mesothelial cells failed to contribute to the intestinal vasculature.

**Figure 6.**
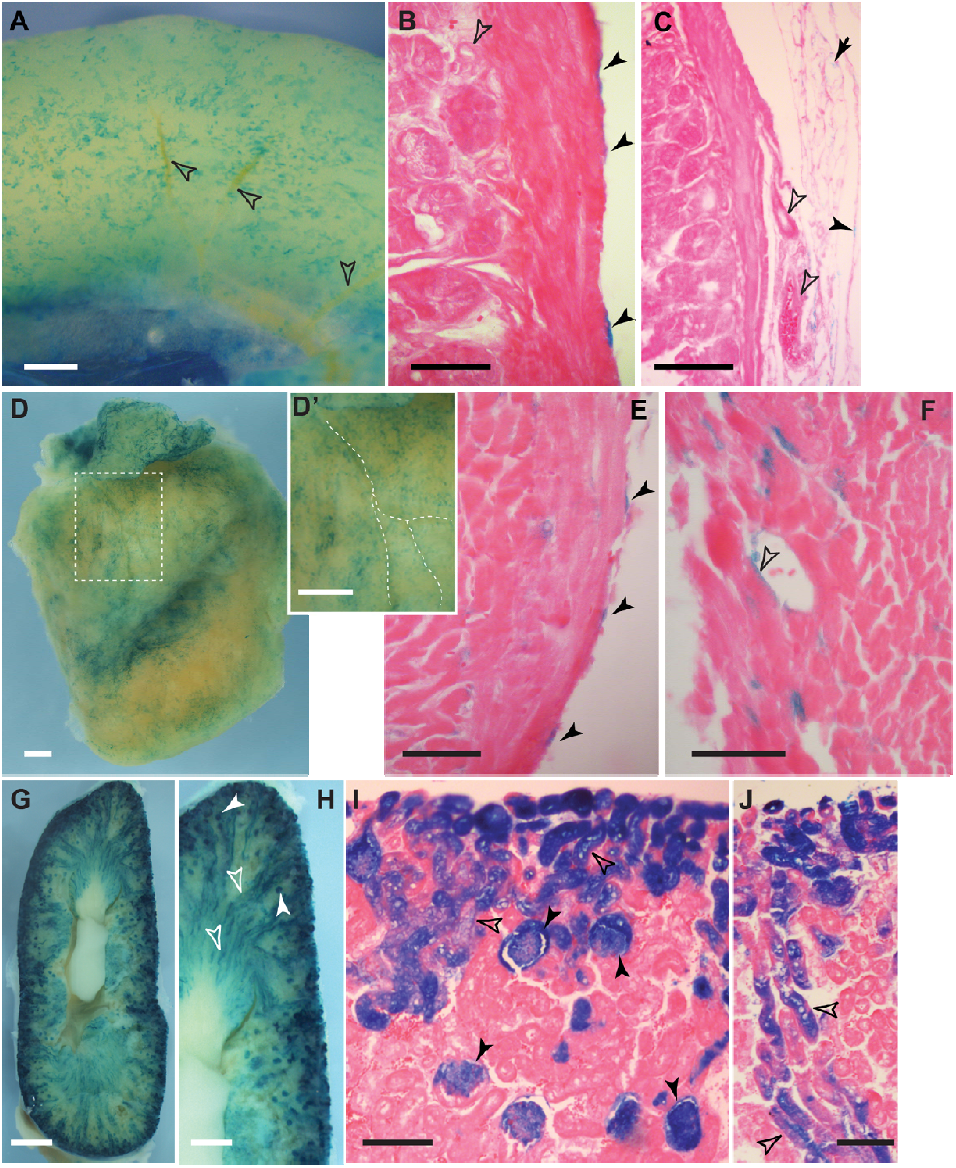
Wt1 lineage tracing in newborn mice. Tamoxifen was given to female animals on days 1 and 4 after giving birth. Intestine (A-C), heart (D-F) and kidneys (G-J) of 7 old Wt1^CreERT2/+^; Rosa26^LacZ/+^ were stained by XGal and analysed for labelled cells. In the intestine, only mesothelial cells were labelled (A, B, filled arrowheads), while there were no labelled cells found around blood vessels (open arrowheads, A-C). In the mesentery, mesothelial (filled arrowhead) and fat cells (arrow) also showed XGal staining (C). The heart (left ventricle and atrium shown, D) was covered with XGal labelled cells localised close to coronary vessels (stippled lines in inset D’). Sections revealed XGal labelling of epicardial cells (filled arrowheads, E), but also of some coronary vessel cells (open arrowhead, F). Labelled cells in the kidneys were abundant throughout (G), predominantly in the glomeruli (H, I, filled arrowheads), but also in nephric tubules (H-J, open arrowheads). Scale bars, 500 μm (A, D, D’, H), 1 mm (G), 50 μm (B, E, F), 100 μm (C, I, J).

In the kidneys, neonate Wt1-expressing cells gave rise to XGal-stained cells in the glomeruli and in nephron tubules (Figure 6G-J). In contrast to lineage tracing in adult kidneys, the contribution of XGal-labelled cells to the nephron tubules was abundant, indicating that Wt1-expressing cells in the neonatal kidneys possess the capacity to give rise to entire nephron structures.

Taken together, our lineage tracing analysis of newborn, juvenile and adult mice demonstrated that Wt1-expressing cells of the peritoneum failed to contribute to the vasculature, or other components of the intestine or body wall musculature besides mesenteric adipose cells. This indicates that in healthy postnatal mice peritoneal mesothelial cells are mostly restricted to self-renewal. Our data showing that Wt1-derived cells in the heart could give rise to coronary vessel cells in newborns and adult mice, suggest that cardiac and peritoneal Wt1-expressing cells may have different capacities to contribute to vasculature in the respective tissue or organ in the postnatal stages.

## Discussion

Here, we have provided novel insight into the postnatal and adult lineage of Wt1-expressing cells in the peritoneum, especially of the visceral and parietal mesothelium, in short- and long-term lineage-traced mice. By utilising conditional lineage tracing of Wt1-expressing cells, our results have revealed that Wt1-expressing mesothelial cells of intestine, mesentery and body wall have mostly a role in maintenance of the peritoneum and fail to contribute to other tissues except visceral adipose tissue.

The mesothelium is a continuous sheet covering the organs housed within the three body cavities, pleural, pericardial and peritoneal, and lining the wall musculature of the cavities. In a previous study, we had used a transgenic mouse line (*Tg(WT1-cre)AG11Dbdr; Gt(ROSA)26Sor/J*; in short Wt1-Cre; Rosa26^LacZ^) composed of a Cre reporter system driven by human Wilms Tumour protein 1 (WT1) regulatory elements ^1^, which had been shown to faithfully recapitulate the Wt1 expression domains in mice ^22^. Using this continuously active Cre reporter system, XGal staining in Wt1-Cre; Rosa26^LacZ^ mice had prominently labelled the vascular smooth muscle surrounding the veins and arteries in the mesentery and those inserting into the intestinal wall ^1^. Based on this finding we concluded that the visceral mesothelium gives rise to these vascular structures during embryonic development. Further, our results led us to hypothesise that there may be a role for the visceral mesothelium in maintaining the vasculature, and possibly other intestinal structures during adult life.

The work presented here, using a conditional tamoxifen-inducible reporter system driven from the endogenous Wt1 locus, demonstrates that the relationship between Wt1 expression and mesothelial lineage in postnatal stages and throughout adulthood is quite simple: The serosa of the peritoneal cavity is predominantly restricted to maintain itself. Our analysis of lineage-traced mice both after short (2-4 weeks) and long (6 months) chase periods combined with clonal analysis demonstrates that self-maintenance as the sole cell fate does not change with time and postnatal mesothelial cells show no other differentiation potential. Therefore, Wt1-expressing cells in the healthy postnatal visceral and parietal peritoneal tissue do not have the capacity to give rise to blood vessel cells, visceral smooth muscle or other non-serosal cells, except for the well-described contribution to visceral adipose cells ^11^. Our Wt1 ablation experiment further revealed that within 10 days after start of tamoxifen dosing, the visceral mesothelium and intestinal wall have maintained their integrity and tissue architecture, suggesting that the Wt1-expressing peritoneal mesothelium is not involved in the homeostasis of the tissues and organs it covers and encases.

Previous pulse-chase studies based on the same tamoxifen-inducible Wt1-driven reporter system as the one used in this report, had shown that the postnatal lung mesothelium makes no contribution to other cells within the lungs ^23^. The same approach had revealed that in the adult liver and the parietal mesothelium of the peritoneal cavity only the Wt1-expressing mesothelium is labelled, indicating self-maintenance of the mesothelium ^24,25^. Together with our own observations, these findings suggest that the Wt1-expressing postnatal peritoneum appears to behave similarly between the lung, liver, intestine as well as parietal peritoneum in that there is no contribution to deeper stromal or parenchymal compartments, other than the progenitor niche of the visceral adipose cells. By contrast, lineage tracing studies after injury in the lungs, liver, heart and peritoneum, including our own, have shown that adult mesothelial cells can be activated and undergo a range of physiological changes including epithelial-mesenchymal transition (EMT) into smooth muscle cells and myofibroblasts, and subsequent contribution to scar formation ^6,25–29^.

Chen and colleagues had reported a minor contribution of the lineage of Wt1-expressing cells to collagen 1a1-expressing submesothelial cells in the visceral and parietal peritoneum of the liver, omentum and body wall ^26^. A recent study confirmed by cytometric sorting the presence of a small population of submesothelial fibroblastic cells that expresses Wt1 together with the mesothelial marker podoplanin and the fibroblast marker PDGFRα, with a possible role in maintaining Gata6 expression in large cavity macrophages ^13^. Here, using the Wt1-based genetic lineage tracing system we have visualized these cells in whole mount preparations of visceral and parietal peritoneum.

In a previous study, the lineage of the mesothelium in a range of internal organs and tissues covered by mesothelial layers had been analysed using a mesothelin-driven LacZ reporter system ^30^. The authors showed that mesothelin-expressing cells in newborn mice contributed to mesenchymal and vascular cells in these organs and tissues, while in adult mice this contribution was reduced. Interestingly, the contribution of mesothelin-expressing cells was particularly prominent to the visceral smooth muscle. This result is conflicting with the findings presented here and elsewhere ^23–25^ using the Wt1^CreERT2/+^; Rosa26^reporter^ system, suggesting that mesothelin-expressing cells undergo a larger range of differentiation processes. It remains unclear how Wt1- and mesothelin-expressing cells differ in their contribution to the maintenance of postnatal and adult tissues and organs covered by mesothelial layers as both markers are expressed in these tissues.

During embryonic development of the heart, Wt1-expressing epicardial cells have been shown to contribute to the mural cells of the coronary vessels as well as fibroblasts ^31–33^. In the adult epicardium Wt1 expression is downregulated ^28^, and lineage tracing studies using the tamoxifen-inducible Wt1-driven lineage system in adult mice have revealed no contribution of epicardial cells to coronary or myocardial cells ^21,34^. Any cells labelled in the vasculature after lineage tracing using the Wt1^CreERT2/+^-based system in adult hearts were suggested to arise from rare Wt1-expessing endothelial cells ^34^. Our data presented here suggest that in newborns but also adult mice, the tamoxifen-inducible Wt1-driven lineage system allows the detection of Wt1-derived cells in the coronary endothelial and mural cells. Based on our results it is not possible to exclude that Wt1-expressing endothelial cells may have contributed to these labelled vascular cells. However, it is striking that Wt1-expressing cells of the postnatal peritoneum failed to provide any contribution to vascular cells. It is possible that the Wt1-expressing cells in the heart (epicardium) and the peritoneum have different capacity to differentiate into mesenchymal or vascular cells in postnatal stages. Alternatively, the absence of Wt1+ endothelial cells in tissue layers underneath the intestinal and parietal peritoneum may account for this difference.

In the kidneys, we observed differences in the contribution to renal tissue between adult and newborn lineage tracing, indicating that there is a larger degree of plasticity present in the newborn kidney, where Wt1-expressing cells contributed to nephron tubules in addition to the glomeruli and parietal epithelial cells of the Bowman’s capsule. These results support the notion that developmental processes of nephron formation and maturation in the newborn kidney in mice is not completed until about postnatal day 3 ^16^.

It is important to point out that the efficiency of cell labelling in lineage tracing systems using the Wt1^CreERT2/+^; Rosa26^mTmG/+^ mouse line has been reported to be between 14.5% and 80% in different laboratories ^26,35^, suggesting that rare lineage and fate changing events may not be detected using this approach ^31^. Therefore, in the current study, we can conclude that XGal- or GFP-positive cells have expressed Wt1 at the time of tamoxifen administration, but there may be some cells that have expressed Wt1 but remained unlabelled and evade the lineage tracing system. The reasons for this variation could be inefficiency of the recombination system or insufficiency of tamoxifen delivery and distribution inside the animals, in combination with variations based on different animal units.

Based on the findings presented here, we propose that Wt1-expressing cells destined to contribute to the vascular smooth muscle component in the intestine and mesentery arise at some timepoint during embryonic development, and not during postnatal stages. Further studies will be needed to elucidate the temporal investment of Wt1-expressing cells in the intestinal and mesenteric vasculature, and the potential role of Wt1 during these processes.

## Methods

### Mice

The following compound mutants were generated for this study by breeding existing mouse lines: Wt1^CreERT2/+^; Rosa26^LacZ/LacZ^ (*Wt1^tm2(cre/ERT2)Wtp^; Gt(ROSA)26Sor/J*) ^10,36^, Wt1^CreERT2/co^; Rosa26^LacZ/LacZ^ (*Wt1^tm2(cre/ERT2)Wtp^; Wt1^tm1.1Ndha^; Gt(ROSA)26Sor/J*) ^37^, Wt1^CreERT2^; Rosa26^mTmG/+^ (*Wt1^tm2(cre/ERT2)Wtp^; Gt(ROSA)26Sor^tm4(ACTB-tdTomato,-EGFP)Luo^/J*) ^38^, Wt1^CreERT2^; Rosa26^LacZ/mTmG^ (*Wt1^tm2(cre/ERT2)Wtp^; Gt(ROSA)26Sor/J; Gt(ROSA)26Sor^tm4(ACTB-tdTomato,-EGFP)Luo^/J*) and Wt1^CreERT2^; Rosa26^Confetti/+^ (*Wt1^tm2(cre/ERT2)Wtp^; Gt(ROSA)26Sor^tm1(CAG-Brainbow2.1)Cle^/J*) ^39,40^. Mice were housed in individually ventilated cages under a 12-hour light/dark cycle, with *ad libitum* access to standard food and water. All animal experiments were performed under a Home Office licence granted under the UK Animals (Scientific Procedures) Act 1986 and were approved by the University of Liverpool AWERB committee. Experiments are reported in line with the ARRIVE guidelines.

### Tamoxifen dosing

Both male and female animals were used in this study. For lineage tracing in adults, 8- to 10-week-old animals were dosed with 100 μg/g body weight of tamoxifen (T5648, SigmaAldrich; 40 mg/ml, in corn oil (C8267, SigmaAldrich)) via oral gavage on 5 consecutive days. Lineage tracing analysis was undertaken at chase times of 2 weeks, after a 10-day wash-out phase, or 5 (Wt1^CreERT2/+^; Rosa26^Confetti/+^) to 6 (Wt1^CreERT2/+^; Rosa26^LacZ/LacZ^) months. For newborn lineage tracing experiments, Wt1^CreERT2/+^; Rosa26^LacZ/LacZ^ male mice were mated with CD1 females (Charles River, Harlow, UK); when a vaginal plug was detected, noon of the day was considered as embryonic day 0.5 (E0.5). Tamoxifen (100 μg/g body weight) was given by oral gavage to the dams at days 1 and 4 after birth, and analysis was performed after a chase of 7 weeks.

For ablation of Wt1 in adults, animals were dosed with tamoxifen (100 μg/g body weight) via oral gavage on 5 consecutive days. Animals were monitored for their well-being, and typically culled at day 10 after the start of the tamoxifen regime.

### XGal staining and histology

Tissues were fixed in 2% paraformaldehyde (PFA)/0.25% glutaraldehyde in phosphate buffered saline (PBS) for between 1 and 1.5 hours, whole-mount XGal staining performed overnight according to standard protocols, followed by overnight post-fixation in 4% PFA (PBS) at 4°C. Histological analysis was performed on post-fixed XGal-stained specimen after dehydration into isopropanol and paraffin embedding. Serial sections (7 μm) were counterstained with Eosin and images taken on a Leica DMRB upright microscope with a digital DFC450 C camera supported by LAS.

### Immunofluorescence on frozen sections

Tissues were fixed in 4% PFA for between 30 to 90 mins, protected in 30% sucrose overnight, placed in Cryomatrix (Thermo Scientific) and snap frozen. Frozen sections were generated at 7 μm on a Thermo Scientific HM525 NX Cryostat. Immunofluorescence analysis was performed following standard protocols ^1^. A bleaching step of 10 min in 3% H_2_O_2_/MetOH was included for embryos or tissues from Wt1^CreERT2^; Rosa26^mTmG/+^ mice in order to remove the tdTomato fluorescence ^25^. The following primary antibodies were used: anti-Wt1 rabbit polyclonal (1:200 to 1:500, clone C-19, sc-192, Santa Cruz), anti-Wt1 mouse monoclonal (1:50, clone 6F-H2, M3561, Dako), anti-SMA mouse monoclonal (1:100 to 1:200, clone 1A4, A2547, Sigma), anti-Pecam/CD31 rat monoclonal (1:50, 550274, Pharmingen), anti-GFP rabbit or goat polyclonal (1:5000, ab6556 or ab6673, Abcam). The anti-SMA antibody was directly labeled using Zenon direct labeling kit (Invitrogen/ThermoScientific) according to manufacturer’s instructions. Secondary antibodies were Alexa fluorophore-coupled (Invitrogen/ThermoScientific) and were used at a dilution of 1:1000. Sections were counterstained with DAPI (D9542, SigmaAldrich) at 1:1000, coverslipped with Fluoro-Gel (with Tris buffer; Electron Microscopy Sciences, USA), and imaged on a Leica DM 2500 upright microscope with a Leica DFC350 FX digital camera and LAS.

### Image analysis

#### Whole mount imaging with and without fluorescence

Imaging of embryos and tissues was performed using a Leica MZ 16F dissecting microscope equipped with a Leica DFC420 C digital camera supported by the Leica Application Suite software package (LAS, version 3 or 4; Leica Microsystems, Germany/Switzerland), and Leica EL6000 fluorescence light source.

Due to uneven tissue geometry, images were taken at different focal levels and subsequently assembled to multilayer composites according to highest focal sharpness.

#### Confetti imaging

Tissues were imaged in form of multilayer Z-stacks with a 3i spinning disk confocal microscope system (Intelligent Imaging Innovations Ltd.) and images subsequently rendered to Z-projection composites using Fiji ImageJ software. For the short- and long-term chase experiments two groups of three animals each were analysed. The small intestine was dissected, cut into 2-3 cm long segments and sliced flat. Four 2 cm long thin tissue segments were randomly chosen and cleaned from feces by multiple PBS washes. Slides were prepared by gluing two layers of 2×22mm coverslips on a standard slide to form an inner rectangular area for tissue placement to prevent leakage during subsequent inverted confocal microscopy. Tissue samples were placed into the space in the correct orientation, PBS added, covered with a standard 22×40mm coverslip and sealed with clear nail polish. Due to the uneven tissue geometry Z-stack images were taken at random where RFP, YFP and CFP cell labelling was identified in close proximity, in some cases any two of the three possible markers. Cells were scored and counted for all three markers in all Z-stacks according to either being a single cell with or without direct contact to a cell of different marker or being in direct contact with another cell(s) of the same marker (designated clones). Statistical analysis was performed by unpaired multiple t-tests with Holm-Šídák multiple comparisons correction using Graphpad Prism 8.4.2.

## Acknowledgements

We acknowledge funding from the Wellcome Trust project grant WT091374MA (BW, NH, TB, TPW) and the Wellcome Trust PhD programme WT102172 (KW, BM, HT), from the MRC project grant MR/M012751/1 (BW, TPW), and self-funded MRes (FM, VF).

We express our thanks to the Biomedical Services Unit at the University of Liverpool, for expert technical support in mouse maintenance and breeding.

## Author contribution statement

TPW and BW designed the experiments, and TPW, HT, FM, VF, BM, KW, TBe performed experiments. TPW and BW interpreted the data and designed the figures. NH and BW contributed to the conception and design of the manuscript. BW wrote the manuscript and all authors were responsible for the approval of the manuscript.

## References

1 Wilm, B., Ipenberg, A., Hastie, N. D., Burch, J. B. E. & Bader, D. M. The serosal mesothelium is a major source of smooth muscle cells of the gut vasculature. Development 132, 5317–5328, doi:10.1242/dev.02141 (2005).

2 Winters, N. I., Thomason, R. T. & Bader, D. M. Identification of a novel developmental mechanism in the generation of mesothelia. Development 139, 2926–2934, doi:10.1242/dev.082396 (2012).

3 Que, J. et al. Mesothelium contributes to vascular smooth muscle and mesenchyme during lung development. Proc Natl Acad Sci U S A 105, 16626–16630, doi:10.1073/pnas.0808649105 (2008).

4 Carmona, R., Cano, E., Mattiotti, A., Gaztambide, J. & Munoz-Chapuli, R. Cells derived from the coelomic epithelium contribute to multiple gastrointestinal tissues in mouse embryos. PloS one 8, e55890, doi:10.1371/journal.pone.0055890 (2013).

5 Colunga, T. et al. Human Pluripotent Stem Cell-Derived Multipotent Vascular Progenitors of the Mesothelium Lineage Have Utility in Tissue Engineering and Repair. Cell reports 26, 2566–2579 e2510, doi:10.1016/j.celrep.2019.02.016 (2019).

6 Karki, S. et al. Wilms’ tumor 1 (Wt1) regulates pleural mesothelial cell plasticity and transition into myofibroblasts in idiopathic pulmonary fibrosis. FASEB journal: official publication of the Federation of American Societies for Experimental Biology 28, 1122–1131, doi: 10.1096/fj.13-236828 (2014).

7 Mutsaers, S. E. et al. Mesothelial cells in tissue repair and fibrosis. Front Pharmacol 6, 113, doi:10.3389/fphar.2015.00113 (2015).

8 Dauleh, S. et al. Characterisation of cultured mesothelial cells derived from the murine adult omentum. PloS one 11, doi:10.1371/journal.pone.0158997 (2016).

9 Wilm, B. & Munoz-Chapuli, R. Tools and Techniques for Wt1-Based Lineage Tracing. Methods in molecular biology (Clifton, N.J.) 1467, 41–59, doi:10.1007/978-1-4939-4023-3_4 (2016).

10 Zhou, B. et al. Epicardial progenitors contribute to the cardiomyocyte lineage in the developing heart. Nature 454, 109–113, doi:10.1038/nature07060 (2008).

11 Chau, Y. Y. et al. Visceral and subcutaneous fat have different origins and evidence supports a mesothelial source. Nature cell biology 16, 367–375, doi:10.1038/ncb2922 (2014).

12 Chau, Y. Y. et al. Acute multiple organ failure in adult mice deleted for the developmental regulator Wt1. PLoS Genet 7, e1002404, doi:10.1371/journal.pgen.1002404 (2011).

13 Buechler, M. B. et al. A Stromal Niche Defined by Expression of the Transcription Factor WT1 Mediates Programming and Homeostasis of Cavity-Resident Macrophages. Immunity 51, 119–130 e115, doi:10.1016/j.immuni.2019.05.010 (2019).

14 Mutsaers, S. E. The mesothelial cell. The international journal of biochemistry & cell biology 36, 9–16 (2004).

15 Boulland, J. L., Lambert, F. M., Zuchner, M., Strom, S. & Glover, J. C. A neonatal mouse spinal cord injury model for assessing post-injury adaptive plasticity and human stem cell integration. PloS one 8, e71701, doi:10.1371/journal.pone.0071701 (2013).

16 Hartman, H. A., Lai, H. L. & Patterson, L. T. Cessation of renal morphogenesis in mice. Developmental biology 310, 379–387, doi:10.1016/j.ydbio.2007.08.021 (2007).

17 Porrello, E. R. & Olson, E. N. A neonatal blueprint for cardiac regeneration. Stem cell research 13, 556–570, doi: 10.1016/j.scr.2014.06.003 (2014).

18 Seely, J. C. A brief review of kidney development, maturation, developmental abnormalities, and drug toxicity: juvenile animal relevancy. Journal of toxicologic pathology 30, 125–133, doi:10.1293/tox.2017-0006 (2017).

19 Cao, J. & Poss, K. D. The epicardium as a hub for heart regeneration. Nature reviews. Cardiology 15, 631–647, doi:10.1038/s41569-018-0046-4 (2018).

20 Porrello, E. R. et al. Transient regenerative potential of the neonatal mouse heart. Science 331, 1078–1080, doi:10.1126/science.1200708 (2011).

21 Quijada, P., Trembley, M. A. & Small, E. M. The Role of the Epicardium During Heart Development and Repair. Circulation research 126, 377–394, doi:10.1161/CIRCRESAHA.119.315857 (2020).

22 Moore, A. W. et al. YAC transgenic analysis reveals Wilms’ tumour 1 gene activity in the proliferating coelomic epithelium, developing diaphragm and limb. Mech Dev 79, 169–184 (1998).

23 von Gise, A. et al. Contribution of Fetal, but Not Adult, Pulmonary Mesothelium to Mesenchymal Lineages in Lung Homeostasis and Fibrosis. American journal of respiratory cell and molecular biology 54, 222–230, doi:10.1165/rcmb.2014-0461OC (2016).

24 Asahina, K., Zhou, B., Pu, W. T. & Tsukamoto, H. Septum transversum-derived mesothelium gives rise to hepatic stellate cells and perivascular mesenchymal cells in developing mouse liver. Hepatology 53, 983–995, doi:10.1002/hep.24119 (2011).

25 Lua, I., Li, Y., Pappoe, L. S. & Asahina, K. Myofibroblastic Conversion and Regeneration of Mesothelial Cells in Peritoneal and Liver Fibrosis. The American journal of pathology 185, 3258–3273, doi:10.1016/j.ajpath.2015.08.009 (2015).

26 Chen, Y. T. et al. Lineage tracing reveals distinctive fates for mesothelial cells and submesothelial fibroblasts during peritoneal injury. J Am Soc Nephrol 25, 2847–2858, doi:10.1681/ASN.2013101079 (2014).

27 Namvar, S. et al. Functional molecules in mesothelial-to-mesenchymal transition revealed by transcriptome analyses. J Pathol 245, 491–501, doi:10.1002/path.5101 (2018).

28 Smart, N. et al. De novo cardiomyocytes from within the activated adult heart after injury. Nature 474, 640–644, doi:10.1038/nature10188 (2011).

29 Kendall, T. J. et al. Embryonic mesothelial-derived hepatic lineage of quiescent and heterogenous scar-orchestrating cells defined but suppressed by WT1. Nat Commun 10, 4688, doi:10.1038/s41467-019-12701-9 (2019).

30 Rinkevich, Y. et al. Identification and prospective isolation of a mesothelial precursor lineage giving rise to smooth muscle cells & fibroblasts for mammalian internal organs, and their vasculature. Nature cell biology 14, 1251–1260, doi:10.1038/ncb2610 (2012).

31 Rudat, C. & Kispert, A. Wt1 and epicardial fate mapping. Circulation research 111, 165–169, doi:10.1161/CIRCRESAHA.112.273946 (2012).

32 Sereti, K. I. et al. Analysis of cardiomyocyte clonal expansion during mouse heart development and injury. Nat Commun 9, 754, doi:10.1038/s41467-018-02891-z (2018).

33 Tian, X. et al. Subepicardial endothelial cells invade the embryonic ventricle wall to form coronary arteries. Cell Res 23, 1075–1090, doi:10.1038/cr.2013.83 (2013).

34 Zhou, B. et al. Adult mouse epicardium modulates myocardial injury by secreting paracrine factors. J Clin Invest 121, 1894–1904, doi:10.1172/JCI45529 (2011).

35 Li, Y., Wang, J. & Asahina, K. Mesothelial cells give rise to hepatic stellate cells and myofibroblasts via mesothelial-mesenchymal transition in liver injury. Proc Natl Acad Sci U S A 110, 2324–2329, doi:10.1073/pnas.1214136110 (2013).

36 Soriano, P. Generalized lacZ expression with the ROSA26 Cre reporter strain. Nat Genet 21, 70–71 (1999).

37 Martinez-Estrada, O. M. et al. Wt1 is required for cardiovascular progenitor cell formation through transcriptional control of Snail and E-cadherin. Nat Genet 42, 89–93, doi:10.1038/ng.494 (2010).

38 Muzumdar, M. D., Tasic, B., Miyamichi, K., Li, L. & Luo, L. A global double-fluorescent Cre reporter mouse. Genesis 45, 593–605, doi:10.1002/dvg.20335 (2007).

39 Livet, J. et al. Transgenic strategies for combinatorial expression of fluorescent proteins in the nervous system. Nature 450, 56–62, doi:10.1038/nature06293 (2007).

40 Snippert, H. J. et al. Intestinal crypt homeostasis results from neutral competition between symmetrically dividing Lgr5 stem cells. Cell 143, 134–144, doi:10.1016/j.cell.2010.09.016 (2010).

