## Supplementary Figures for "Restricted differentiative capacity of Wt1-expressing peritoneal mesothelium in postnatal and adult mice"

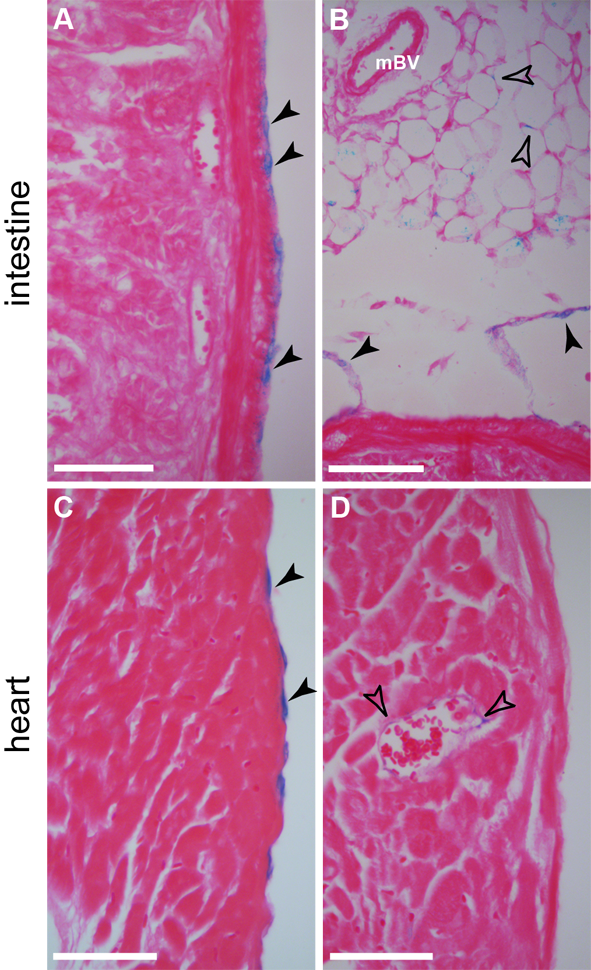

**Supplement Figure S1: Histological analysis of mesothelial contribution to adult intestine and heart after lineage tracing of Wt1-expressing cells.** Adult Wt1^CreERT2/+^; Rosa26^LacZ/LacZ^ mice were analysed 2-4 weeks after Tamoxifen administration. A, B. LacZ-positive cells in eosin counterstained paraffin sections of intestine (A) and mesentery (B) were detected in the serosal mesothelium (solid arrowheads) as well as in the mesenteric fat (hollow arrowheads); mBV, mesenteric blood vessel. Open arrowhead points towards mesenteric blood vessel. C, D. Eosin counterstained sections through ventricular wall of the heart revealed LacZ-positive cells in the epicardium (arrowheads, C) and in the coronary vessels (hollow arrowheads, F). Scale bars, 50 µm (A, C, D), 100 µm (B).

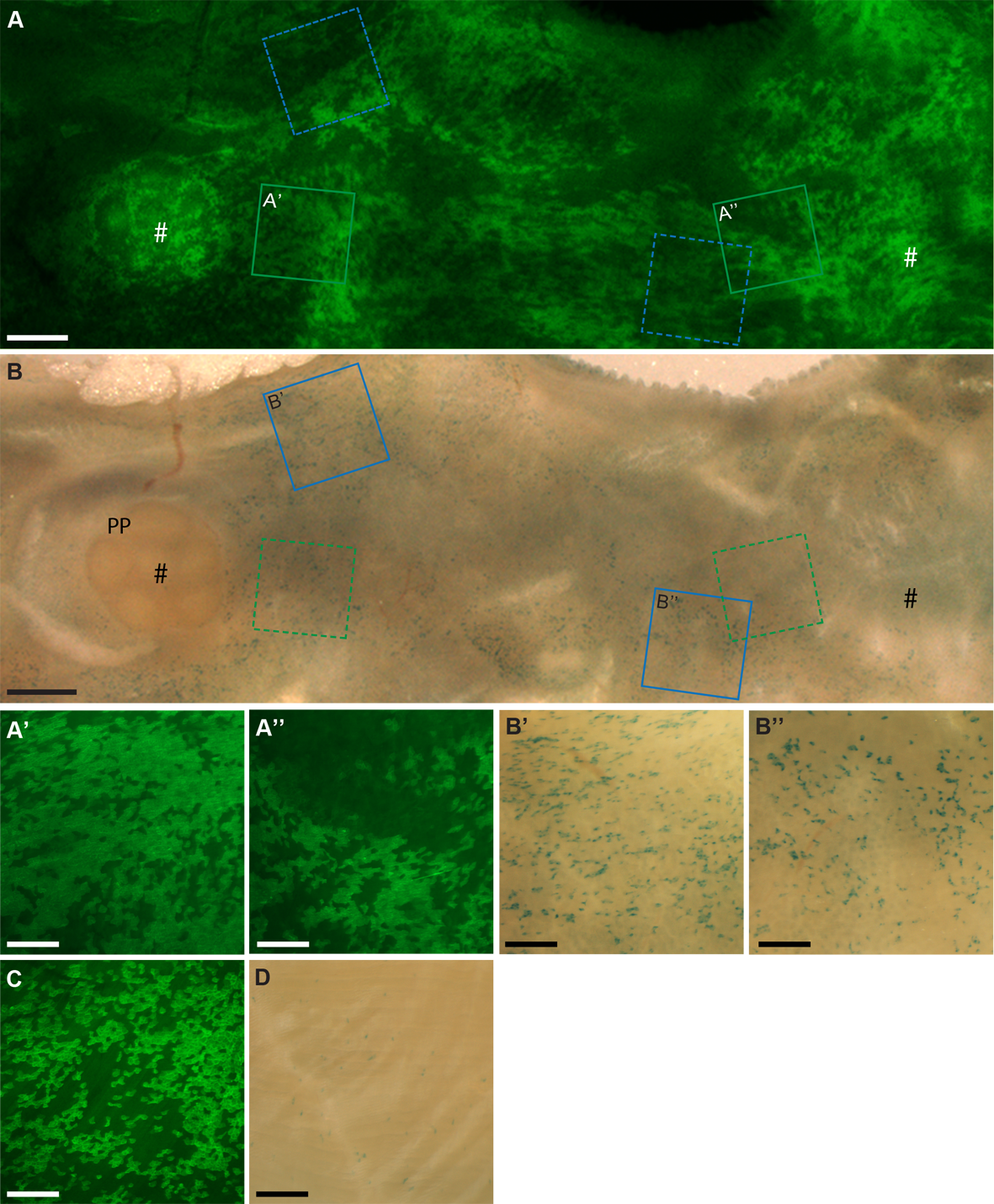

**Supplementary Figure S2: Comparison of the distribution of peritoneal GFP- and LacZ-expressing cells in the intestine and body wall muscle after lineage tracing in Wt1^CreERT2/+^; Rosa26^LacZ/mTmG^ reporter mice.** A, B. A segment of small intestine was dissected and carefully stretched out on a wax plate using fine insect pins and the profile of the GFP-expressing cells captured (A). The stretched-out tissue segment was then fixed, removed from the wax plate, XGal-stained and reattached to the same wax plate (using matching pin wholes) for capturing of the profile of LacZ-positive cells (B). In direct comparison many more GFP-positive cells overall were detected than LacZ-positive cells. Also, LacZ-free areas showed much more frequently GFP-positive cells (#) than vice versa. Specific regions of the intestinal tissue were imaged at higher magnification to demonstrate irregular coverage of GFP-expressing (A’, A’’) or LacZ-expressing cells (B’, B’’; highlighted in A and B, respectively). C, D. Coverage of GFP-positive cells in the parietal peritoneum (of the body wall muscle layer; same animal) was similarly dense while sparse for LacZ-positive cells. Peyer’s patch (PP); scale bars, 1mm (A, B), 300 µm (A’, A’’, B’, B’’, C, D).

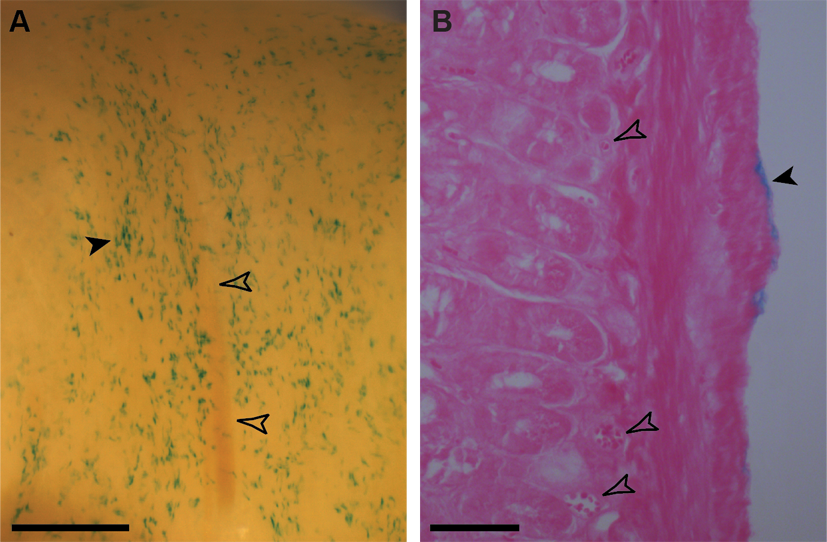

**Supplementary Figure S3: Long-term lineage tracing of Wt1-expressing cells in adult mouse intestine.** Adult Wt1^CreERT2/+^; Rosa26^LacZ/LacZ^ mice were analysed 2 and 6 months after Tamoxifen administration (n = 6 in both groups). A. Whole mount analysis revealed no difference in distribution pattern and patchiness of LacZ expressing cells in the serosal mesothelium in either group (long chase shown only; filled arrowhead pointing to group of neighbouring cells, open arrowheads to blood vessel). B. Eosin counterstained paraffin sections of the same intestinal segment as shown in A, demonstrated LacZ expressing cells in the mesothelium overlying the muscularis of the intestinal wall (filled arrowhead), but no other tissues including the vasculature (open arrowheads). Scale bars, 400 µm (A) and 50 µm (B).
